# Behavioral context affects social signal representations within single primate prefrontal cortex neurons

**DOI:** 10.1101/2021.11.01.466818

**Authors:** Vladimir Jovanovic, Adam R. Fishbein, Lisa de la Mothe, Kuo-Fen Lee, Cory T. Miller

## Abstract

We tested whether social signal processing in more traditional, head-restrained contexts is representative of the putative natural analog – social communication – by comparing responses to vocalizations within individual neurons in marmoset prefrontal cortex (PFC) across a series of behavioral contexts ranging from traditional to naturalistic. Although vocalization responsive neurons were evident in all contexts, cross-context consistency was notably limited. A response to these social signals when subjects were head-restrained was not predictive of a comparable neural response to the identical vocalizations during natural communication, even within the same neuron. Neural activity at the population level followed a similar pattern, as PFC activity could be reliably decoded for the context in which vocalizations were heard. This suggests that neural representations of social signals in primate PFC are not static, but highly flexible and likely reflect how nuances of the dynamic behavioral contexts affect the perception of these signals and what they communicate.

## Introduction

Communication is an inherently interactive social process characterized by the active exchange of signals between conspecifics (Guilford and Dawkins, 1991). Yet because of practical constraints, experiments seeking to explicate the neural basis of social communication in the primate brain have traditionally employed paradigms in which social signals – such as faces and vocalizations – are presented as static stimuli completely divorced from the dynamic behavioral contexts in which they naturally occur. While these passive-viewing/listening approaches have been notably prolific at revealing integrated networks for both face and voice processing in primate temporal and frontal cortex (Freiwald et al., 2016; Gifford et al., 2005; Hung et al., 2015; Perrodin et al., 2011; Petkov et al., 2008; Romanski et al., 2005; Schaeffer et al., 2020; Tsao et al., 2006; Tsao and Livingstone, 2008; Tsao et al., 2008), fundamental questions remain about the role that these neurons play during active communication. Indeed, McMahon and colleagues (McMahon et al., 2015) found that although neurons in the AF face patch exhibited classic selectivity to face stimuli when subjects passively viewed static images of faces, the responses of the same individual neurons were primarily driven by a myriad of different properties – including social proximity and movement – when subjects simply watched videos of monkeys engaged in natural social interactions. Such findings raise a critical question of whether neural responses to passively presented social stimuli in conventional primate head-restrained paradigms are representative of how the same individual neurons (or populations) will respond when subjects are actively participating in natural communication exchanges. Here we sought to test this critical issue by directly comparing the responses of individual, single neurons in marmoset PFC to conspecific vocalizations in a series of behavioral contexts – ranging from a more traditional, head-restrained paradigm to freely-moving monkeys engaged in interactive natural communication (**Figure 1A**). We hypothesized that if the activity of neurons in a more traditional context was representative of natural communication, similar patterns of neural activity would be evident within individual units and/or at a population level across context. If, however, neural responses to vocalizations in one behavioral context poorly predicted another, it would demonstrate a more dynamic system in which behavioral context affects the perception of social signals and the related neural representations in primate PFC. Such a result would also highlight the caveats of using traditional primate head-restrained paradigms to elucidate the neural basis of natural primate social behaviors.

**Figure 1.**
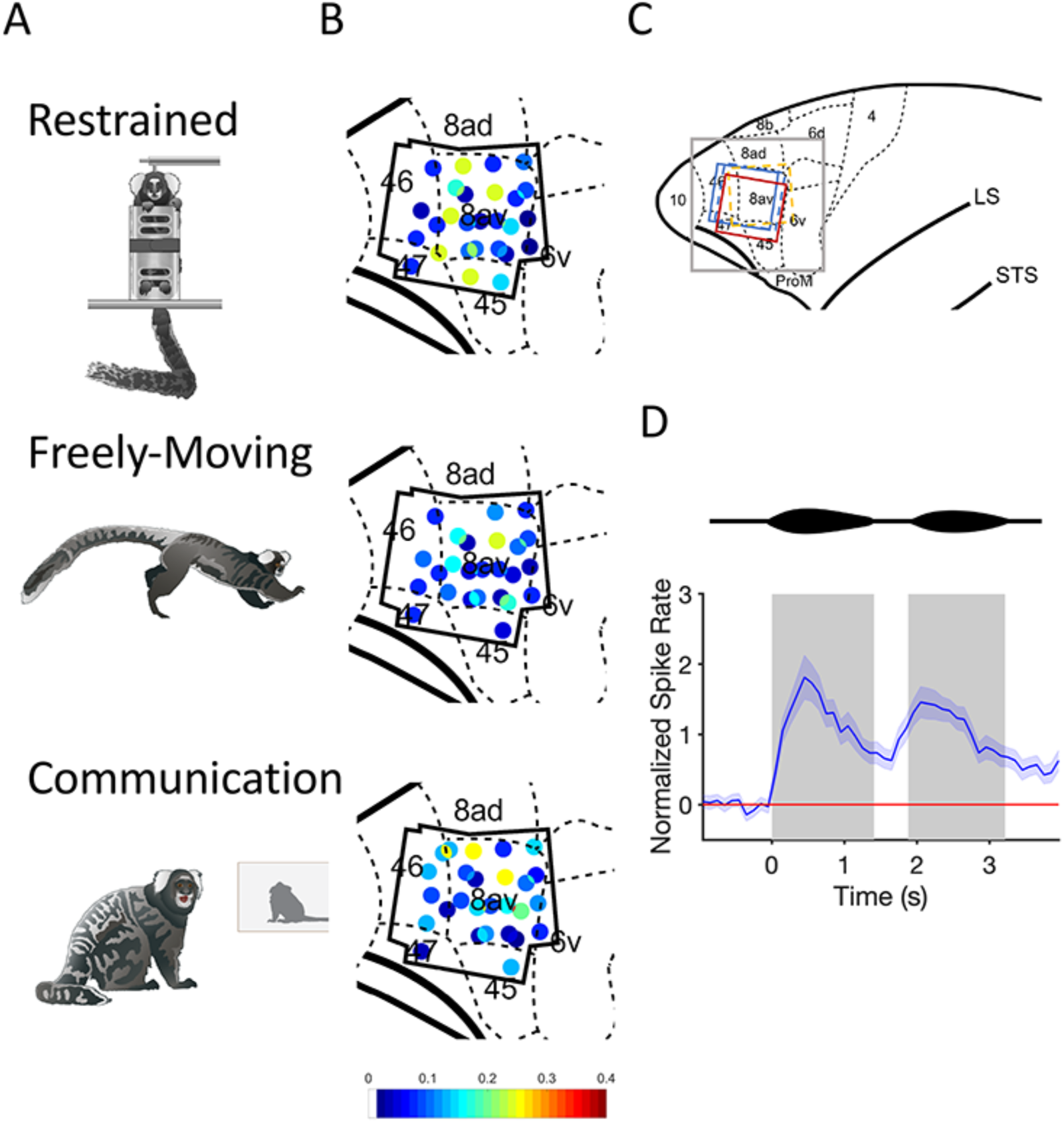
(A) Schematic drawings of the three behavioral contexts. (B) Distribution of phee responsive neurons in marmoset prefrontal cortex. Black polygonal shape represents the outline of the four electrode arrays across three marmosets. Each dot represents one electrode channel and its ratio (indicated by the color bar) of phee responsive neurons relative to all single neurons recorded at that location. A separate map is shown for each behavioral context. (C) Anatomical map of the frontal cortex with labeled brain areas. Gray square represents the zoomed in portion of PFC depicted in B. Other colored squares represent the position and orientation of the four electrode arrays. Dashed lines represent right hemisphere implants, and solid represent left hemisphere. (D) A schematic depiction of a two-pulsed marmoset phee call is shown above a normalized PSTH with 95% Confidence Interval for all phee-responsive neurons across all behavioral contexts. Gray boxes indicate the average duration of phee pulses.

## Results

### Vocalization Responses in Marmoset PFC

Here we recorded the activity of 388 single neurons in the PFC of three marmoset monkeys (*Callithrix jacchus*) in response to vocalizations in three behavioral contexts – Restrained, Freely-Moving and Communication (Figure 1A). In the ‘Restrained’ condition, head-restrained marmosets were seated in a primate chair and presented a battery of acoustic stimuli at a fixed inter-stimulus interval. The ‘Freely-Moving’ condition had identical stimulus presentation, but the animals were able to move unrestrained within the test box. During the ‘Communication’ condition, freely-moving marmosets engaged with a Virtual Marmoset (VM) in their natural antiphonal conversations using interactive playback software implemented in several previous behavioral and neurophysiological experiments (Miller et al., 2009; Miller et al., 2015; Miller and Wang, 2006; Nummela et al., 2017). Because marmosets only produce phee calls during these natural conversations, we focused analyses on neural responses to hearing this call type across all behavioral contexts. In order to test for any contextual differences in neural activity, identical phee calls produced by a single caller were presented to subjects in all three contexts within each daily test session. At sites across all areas of PFC examined here (i.e. 8av, 45, 46 and 47, Figure 1B,C) and in all three contexts, we found neurons that exhibited significant changes in neural activity from baseline in response to hearing marmoset phee calls.

### Within-Neuron Comparisons

This study was designed to directly test whether two key behavioral characteristics that differ between traditional and naturalistic studies of communication significantly affected PFC responses to vocalizations within individual neurons - subjects’ mobility (head-restrained or freely-moving) and stimulus presentation (consistent timing interval or interactive). As a result, we focused the next set of analyses on the 247 single PFC neurons that maintained consistent isolation across all three behavioral contexts. We hypothesized that if subjects’ mobility was a key factor modulating vocalization responsiveness within PFC neurons, neural activity in the Restrained context would be distinct from the other two. By contrast, if stimulus presentation significantly affected responses to vocalizations, neural responsiveness in the consistent interval conditions (Restrained and Freely-Moving) would be similar to each other and differ from the interactive context (Communication). If, however, a broader suite of contextual features affects vocalization-responsivity in primate PFC neurons, we would expect little consistency across the behavioral contexts.

Overall, data were consistent with the latter hypothesis. Although neurons exhibited robust stimulus-driven responses to marmoset phee calls across the behavioral contexts (Figure 1D), we observed a remarkable lack of cross-context consistency in the pattern of vocalization responsiveness in this population. Significant neural responses to phee calls in one behavioral context poorly predicted a comparable response in another. Figure 2A shows the percentage of units found exhibiting a response to phee calls within and between each individual context. Overall, 170 neurons exhibited a significant change in activity in response to phee calls. Of these neurons, 29 units were responsive only in Restrained, 34 only in Freely-Moving, 38 only in Communication, 13 in both Restrained and Communication, 15 in both Restrained and Freely-Moving, 15 in both Freely and Communication, and 26 neurons exhibited a significant change in activity in all three behavioral contexts – i.e. Restrained, Freely-Moving, Communication – and described as RFC units in subsequent analyses. Overall, the probability that a vocalization responsive neuron in one behavioral context would exhibit the same response in another context was only 16.3% (SD = 0.418). This pattern of acoustic stimulus responsivity and contextual heterogeneity to was not limited to phee calls, as it was also evident when comparing neural responses to a broader corpus of marmoset vocalizations and noise stimuli between the Restrained and Freely-Moving contexts (Figure S1).

**Figure 2.**
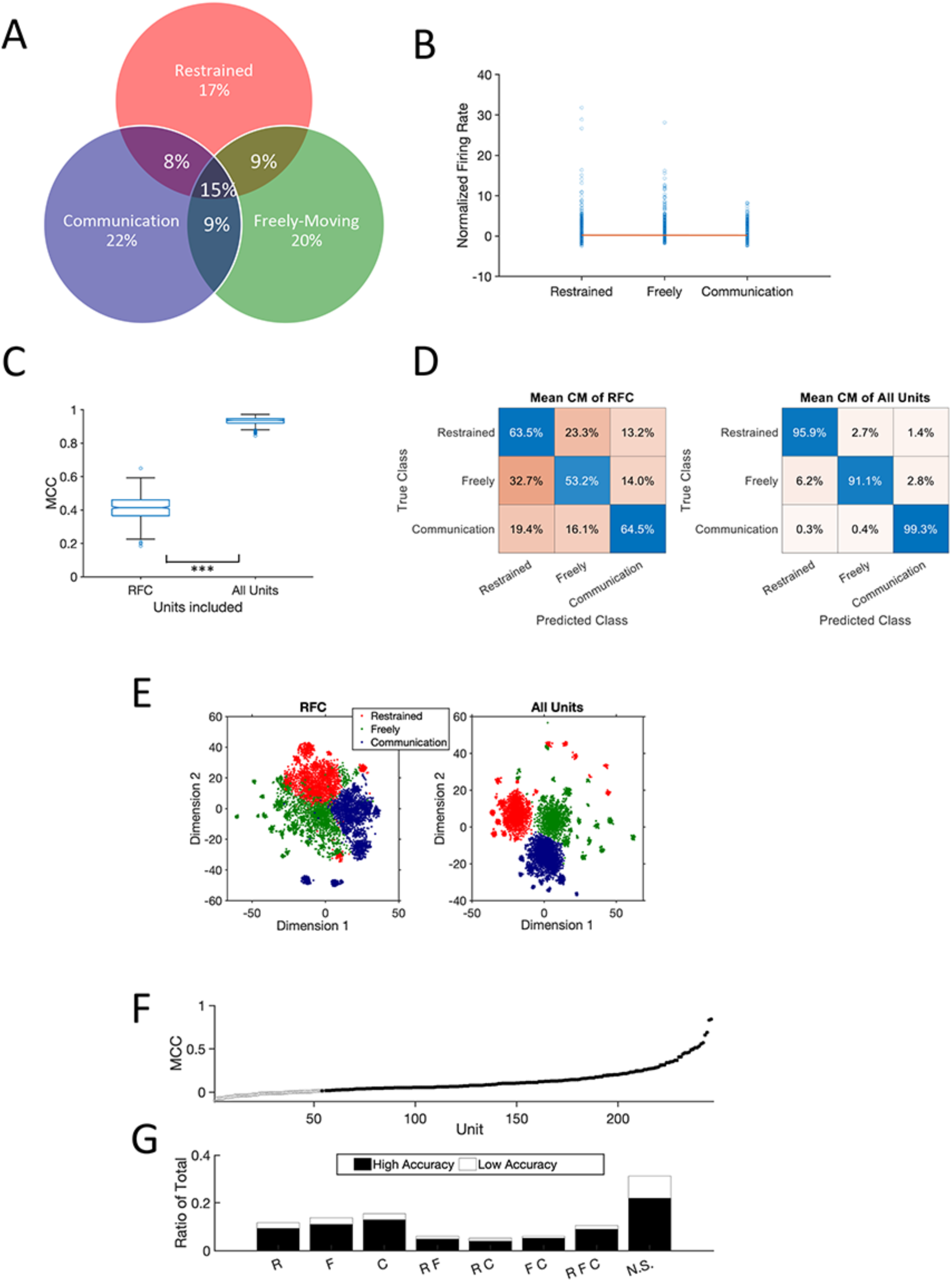
(A) Venn Diagram depicts the heterogeneous responses to marmoset phee calls in the 247 well-isolated single units maintained across all three contexts for a daily test session. The values are the percentages of all significantly responding units (170 units) for each combination of contexts. (B) Normalized firing rate to phee calls in each of the three behavioral contexts. Single blue circles represent each individual phee-responsive neuron. The red line was constructed from the slope and intercept of the linear mixed effect model that did not show a significant effect of context on firing rate. (C) Performance of neural decoder for RFC units and All Units in this population. Simulations performed above chance for RFC units, but this performance was significantly less than when All Units were included in the analysis. (D) Mean confusion matrix across 500 simulations decoders tested with only RFC units (left) and All Units (right). Percentages are row-normalized, showing how all the true classes were binned across the predicted classes. Each row sums to 100%. (E) t-SNE plot (t-Distributed Stochastic Neighbor Embedding) created by inputting the median performing simulation’s PCA plot. Note that RFC has much more overlap, and that a k-means clustering of the three classes would perform poorly compared to All Units. (F) Individual unit performance in decoding neural response into the three behavioral contexts. 95% confidence intervals are shown but too small to be visible. Filled in points represent units that had significantly higher response above chance in accuracy (p = 0.001). (G) Distribution of units that had significantly higher accuracy above chance (black) compared to below chance (white) binned across the category types from panel A. N.S. refers to units that lacked any significant response to phee calls.

Several factors other than the dynamic nature of behavioral contexts could explain the pattern of results here. One possibility is that differences in attention and or arousal might drive a linearly additive effect on neural responses across behavioral contexts (Restrained < Freely < Communication). However, analysis of overall firing rate indicated that vocalization-driven activity was remarkably similar across the contexts (Figure 2B), and a linear mixed effect model failed to find evidence that firing rate changed linearly between the contexts (F(1, 2.015) = 4.017, p = 0.182, n = 29164 observations). A second possibility is that PFC response heterogeneity could result from spatial selectivity due to head-direction. We systematically changed subjects’ spatial position relative to the speaker in the Restrained chair and measured the relative angle of the head to the speaker in both the Freely-Moving and Communication contexts during stimulus presentations in a subset of neurons. In each case, the vast majority of neurons tested did not exhibit any spatial selectivity (Figure S2A), a result consistent with previous experiments in primate PFC that likewise failed to find spatial modulation of auditory responses (Cohen et al., 2009). Lastly, because phee stimulus timing differed between the Restrained/Freely-Moving contexts and Communication, it is possible that differences in the inter-stimulus interval inadvertently affected neural responses. We examined the ratio in the firing rate between two consecutive phee calls in each of the three behavioral contexts based on the time interval between each pair of consecutive phee stimuli to test whether calls with a shorter inter-stimulus interval would be more likely to exhibit similar firing rates. Analyses failed to find any evidence of such an effect, as the standard deviation in firing rate was similar across all behavioral contexts regardless of the interval between the phee call stimuli (Figure S2B).

Evidence showed that the heterogeneity of PFC responses was also evident at the population level. We applied a decoder to test if patterns of neural responses differed by context across the population. We independently tested two sets of neurons – RFC Units (n=26) and All Units (n=247) - as inputs across 500 simulations and used the Matthew’s Correlation Coefficient (MCC, +1 perfect prediction, 0 for random, -1 total disagreement) to evaluate performance. We hypothesized that the decoder for all units would be highly accurate in predicting the context and would outperform the RFC units, consistent with a mixed-selectivity population level coding scheme (Bernardi et al., 2020; Blackman et al., 2016; Fusi et al., 2016; Parthasarathy et al., 2017; Rigotti et al., 2013). Indeed, we observed that All Units exceeded 90% accuracy in all contexts and significantly outperformed RFC Units (Wilcoxon signed rank test, n = 500, signed rank = 0, p < 0.001), despite the fact that only the latter population comprised vocalization-responsive neurons in each of the three behavioral contexts (Figure 2C). The confusion matrices in Figure 2D illustrate other notable decoding differences between these populations. First, the decoder for All Units had an overall substantial increase in accuracy for all contexts relative to RFC Units. Second, though more pronounced for the RFC Units, both decoders revealed higher false positives between the Restrained and Freely-Moving contexts. The overlap between these populations is also evident through dimensionality reduction in a principal component analysis (Figure 2E). Although neural activity in these two contexts was accurately decoded at over 90% for All Units, it does suggest at least a modicum of similarity considering that the decoding accuracy for the Communication context was nearly perfect (99.3%).

Given the notably improved accuracy of the decoder that included All Units, we tested performance at the individual unit level. For all 247 units, we ran 500 simulations and calculated their 99.99% confidence interval to determine which ones were above chance in overall accuracy performance. We found that 193 units (78%) had significantly better performance than chance (Figure 2F). Interestingly, the proportion of high accuracy units within each class of phee-responsive neurons was roughly similar across the population (Figure 2G) suggesting that the context-dependence of stimulus-driven activity may not be a significant factor for decoding.

### Active vs Passive Listening during ‘Communication’

We have already shown that behavioral context can be decoded from PFC neural activity. An additional question is whether task demands such as active vs. passive engagement with stimuli can be similarly decoded within a behavioral context. Indeed, task demands are known to affect PFC activity in primates (Parthasarathy et al., 2017), including for tasks in which subjects are trained to respond to vocalizations (Cohen et al., 2009; Hwang and Romanski, 2015; Plakke et al., 2013). Here, we took advantage of the fact that the Communication context comprises instances when marmosets passively listen to a conspecific call, but do not produce a vocal response (Independent), and occasions when marmosets produce a vocalization response upon hearing a conspecific (Antiphonal) (Nummela et al., 2017). We hypothesized that if the difference between passive-listening and active behavior significantly affected neural activity, it would be easily decoded. Using the same type of decoder described above, however, we found performance across all the units was notably low in MCC score with a mean of 0.373 (SD = 0.160) and in accuracy (mean = 0.630, SD = 0.0854). These results suggest that PFC activity during the stimulus-presentation period did not differ within context based on task engagement.

This analysis focused specifically on stimulus-driven activity, but previous results indicated that inclusion of neural activity in the period of time immediately preceding and following a vocalization revealed the presumptive social state of marmoset frontal cortex and reliably predicted the likelihood of a vocalization response (Nummela et al., 2017). We applied that same approach to the dataset here and found a significant increase in the decoder’s ability to distinguish between phee calls heard in the Independent and Antiphonal contexts over the stimulus-driven analysis above (Wilcoxon signed rank test, n = 500, signed rank = 35, p < 0.001; Figure 3B). The confusion matrices shown in Figure 3C further illustrate the difference in decoding accuracy between these analyses, with significantly improved performance when including the pre- and post-stimulus firing rate. Lastly, we analyzed the individual performance of these units to decode Antiphonal and Independent events within the ‘Communication’ context for this latter state-responsive analysis. We found 55 of 94 units had a significantly higher accuracy than chance with confidence interval at 99.9% (Figure 3D). Even PFC neurons without a significant stimulus-driven response to phee calls in the ‘Communication’ context provided meaningful information to distinguish between these two behavioral contexts (Figure 3E), showing the importance of such neural activity during natural social interactions.

**Figure 3.**
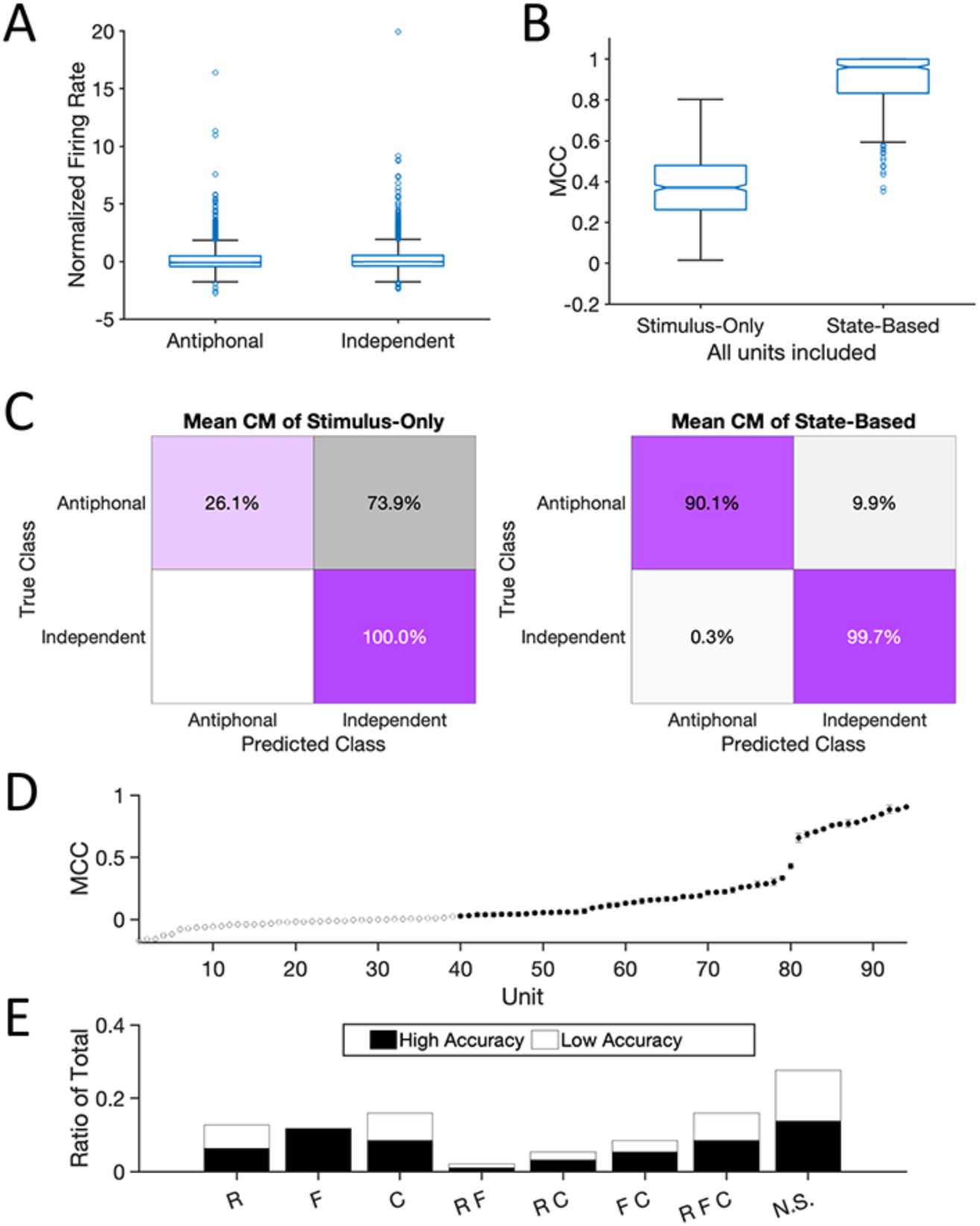
(A) Box plots showing the normalized firing rate of neurons exhibiting a stimulus-driven response in the ‘Antiphonal’ and ‘Independent’ phee call contexts within ‘Communication’. Single blue circles represent individual neurons. (B) Outcome of the decoder applied to these two categories (Antiphonal and Independent) in the Communication context using units that had at least 5 trials in both categories for Communication context. ‘Stimulus-Only’ refers to the decoder that included firing rate from time bins only during the phee call stimulus. ‘State-Based’ refers to the coder that included the phee stimulus as well as pre and post-phee stimulus bins. (C) Mean confusion matrix for decoder performance for Stimulus-Only (left) and State-Based (right). Blank space means no Independent categories were misclassified as Antiphonal in the Stimulus-Only decoder. (D) Individual unit performance of the 94 units in the State Based decoder. While 39 units had accuracies at or below chance, 55 had accuracies above with 99.9% confidence. 95% confidence interval bounds are plotted but are too small to be visible. (E) Distribution of High (Black) and Low (White) accuracy units from the State-Based decoder is shown for each class of neural response category times from Figure 2A.

## Discussion

Neurophysiological studies of social signal processing have typically relied on passive, head-restrained paradigms, but here we found that PFC activity in this context was not predictive of the putative natural analog – communication. Instead, we found that the context in which vocal signals are heard profoundly affects their representation in primate PFC. Although vocalization responsive neurons were evident in each of the three contexts tested here, the pattern of responses was highly heterogeneous both within and between neurons. Despite the fact that vocalization-responsive neurons were evident in each of the three behavioral contexts, a decoder could almost perfectly classify the context in which the calls were heard from PFC activity, suggesting dynamic population-level differences in how these social signals are represented. In other words, it is not simply that different neurons are responsive to vocalizations in different contexts but that these responses are integrated at a population level for a consistent representation. Rather, the results here are consistent with a mixed-selectivity mechanism, in which task-relevant information is distributed across a neural population to support flexible behaviors (Bernardi et al., 2020; Blackman et al., 2016; Fusi et al., 2016; Parthasarathy et al., 2017; Rigotti et al., 2013), such as natural primate communication. By holding the stimulus – phee calls - constant and manipulating the context in which they were heard, our findings reflect the fact that the perception of primate social signals is not static, but highly affected by the contextual nuances in which they occur (Cheney and Seyfarth, 2018). Our data indicate that the neural representations of social signals in traditional head-restrained paradigms is neither representative of natural communication nor reflects a baseline state upon which behavioral demands modulate activity. Rather, the behavioral contexts in which vocalizations occur are parallel, with each having distinct effects on their neural representation in primate PFC.

Contextual differences in the neural representations of vocal signals likely reflect how contextual variables affect the perception of social signals. Marmosets only perceive conspecifics to be communicative partners if they abide species-specific social rules during vocal interactions (Miller et al., 2009; Toarmino et al., 2017). Therefore, while marmosets did hear phee calls in each context, those calls were likely only perceived as being produced by a communicative partner in the Communication context. By contrast, phee call stimuli in both the Restrained and Freely-Moving contexts were not broadcast in accordance with the temporal dynamics of marmoset conversations, but rather were randomized with other vocalizations and noise stimuli at a regular temporal interval. Neural responses at the individual unit and population levels were not uniform between these contexts, nor linearly additive (Figure 2). While animals frequently eavesdrop on conspecific vocal interactions (McGregor, 2005; Mennill et al., 2002), the timing of phee presentations here was artificial and unrelated to the natural statistics of marmoset communication. These results suggest that the properties distinguishing behavioral contexts and their impact on social signal perception likely exist in a high-dimensional space that mirror the dynamics of primate sociality – and includes computational flexibility to adapt to contexts that do not occur naturally.

Interestingly, two behavioral conditions that could not be accurately decoded from stimulus-driven activity were within natural communication. Specifically, during the Communication context the decoder was unable to correctly classify neural responses to vocalizations heard in two behaviorally distinct events typical of natural marmoset communication - phee calls that elicited a vocal response from conspecifics (Antiphonal) and those that did not (Independent). Although subjects may not have produced a vocal response each time they heard a conspecific call, they likely perceived the conspecific VM as a willing communicative partner because that individual had abided social rules throughout the session. Stimulus-driven neural responses, however, are not the only signatures of neural activity during natural marmoset vocal interactions (Nummela et al., 2017). A second decoder that included neural activity in the period before and after each phee call stimulus was able to almost perfectly classify these natural behavioral contexts (Figure 3). Consistent with our previous finding (Nummela et al., 2017), this suggests that the presumptive social state of PFC neurons underlies natural primate social interactions, and may function as a complementary mechanism to mixed-selectivity for primate social brain functions.

Communication is social behavior for which signal processing is one component, rather than a system in which a signaling processing system forms the base on top of which behaviors are built. The current study focused solely on PFC but there are reasons to think that the effects of social context on primate neocortical function are more widespread (Ainsworth et al., 2021; Cléry et al., 2021; Sliwa and Freiwald, 2017). Indeed, merely the presence of conspecifics can affect primate decision-making (Chang et al., 2013; Chang et al., 2011), while simply watching conspecific social interactions as a third-party observer has been shown to change the response properties of classically identified face cells to be driven by a myriad of social factors (McMahon et al., 2015). Coordinated social and vocal interactions drive brain-to-brain coupling in the frontal and temporal cortex of humans and other mammals that may be a unique neural signature to coordinated social interactions (Stephens et al., 2012; Zhang and Yartsev, 2019). By demonstrating - for the first time – that the paradigms routinely employed to explicate the neural basis of communication in the primate brain do not faithfully reflect PFC activity during the analogous natural social behavior, we open the door to further questions about modeling the neural dynamics of active, freely-moving behaviors. As research pushes further towards examining the longitudinal and idiosyncratic facets of natural social interactions that measures behavior at a temporal resolution more closely matching the brain (Calhoun et al., 2019; Pereira et al., 2019; Pereira et al., 2020), we are poised to leverage the power of these technologies to gain deep insight into natural primate social brain functions.

## Acknowledgements

This work was supported by grants from the NIH to CTM (2R01 DC012087). All experiments were approved by the UCSD IACUC (S09147).

## STAR Methods

### Subjects

Three adult common marmosets (Callithrix jacchus) were used for this experiment. H01 and E01 were female while H02 was male. All subjects were at least 1.5 years old at time of implant. E01 had bilateral arrays implanted. H01 had a right hemisphere PFC implant, while H02 had a left hemisphere. All animals were group housed, and experiments were performed in the Cortical Systems and Behavior Laboratory at University of California San Diego (UCSD). All experiments were approved by the UCSD Institutional Animal Care and Use Committee.

### Behavioral Contexts

Subjects were tested on three behavioral contexts in each session: ‘Restrained’, ‘Freely-Moving’ and ‘Communication’. The order of these contexts was randomized each session and counterbalanced between subjects. Within each recording session, subjects were presented with the same set of phee calls produced by a single caller from the UCSD colony and recorded previously following standard vocal recording procedures (Miller and Wang, 2006). A total of 10 adult marmoset were used to generate the phee call stimulus sets and were randomly selected for each test session.

- <u>Restrained</u>. Marmosets were head-restrained in a standard chair used in previous research (Mitchell et al., 2014; Nummela et al., 2019) and a series of acoustic stimuli with a 1 s inter-stimulus interval that comprised phee calls and 1 s noise was broadcast. The stimulus sets broadcast to subjects H01 and H02 also included reversed phee calls, twitters, and reversed twitters. Stimuli were organized into four blocks and the stimulus order was randomized. Thirty exemplars of each stimulus type (i.e. phees, noise, etc) were broadcast in every test session.
- <u>Freely-Moving</u>. The identical stimulus presentation protocol as described for ‘Restrained’ was used in this context. The only difference was that the animals were freely-moving in the test box rather than head-restrained in a primate chair.
- <u>Communication</u>. Similar to the Freely-Moving context, subjects here were able to move freely around the test box. Rather than broadcast the same stimuli as the previous two conditions, only phee calls were presented using our interactive playback paradigm. Here, marmosets engaged with a computer-generated Virtual Marmoset (VM) designed to broadcast phee calls in response to subjects’ calls and simulate natural vocal interactions. The identical paradigm has been used in several previous behavioral and neurophysiological studies of marmosets in our lab (Miller and Thomas, 2012; Miller et al., 2015; Nummela et al., 2017; Toarmino et al., 2017). Briefly, the software is designed to detect phee calls produced by subjects online. Whenever subjects produce a phee call, the VM emits a phee call within 2-4 s in response. If subjects do not emit a phee call for more than 45-90 s, the VM will also broadcast a phee call. Phee calls produced by subjects within 10 s of a VM phee call are classified as an ‘antiphonal’ response. Calls produced outside of that time period are classified as ‘spontaneous’ phee calls.

### Test Procedures

All recording sessions took place in a Radio-Frequency shielding room (ETS-Lindgren) in a 4 × 3 m room. Subjects were placed in a clear acrylic test box with a plastic mesh on the front side (32 × 18 × 46 cm) or standard primate chair positioned on a table on one side of the room. A single speaker (Polk Audio TSi100, frequency range 40-22,000 Hz) was placed 2.5 m away from the test box on the opposite side of the room. A black cloth occluder was positioned equidistant between the table and speaker to eliminate subjects’ ability to see the speaker. One microphone was placed in front of the subject and speaker each (Sennheiser, model ME-66). The speaker broadcast acoustic stimuli at an approximate 80-90 db SPL measured 1 m from the speaker. Subject and speaker calls were recorded simultaneously with the neurophysiological data on a data acquisition card (NI PCI-6254).

Subjects H01 and H02 also participated in a separate ‘Orientation’ test condition to determine whether neurons in marmoset PFC were spatially selective. Here subjects were positioned in a chair and head-restrained, similarly to the Restrained context, and tested in the same test room but the relative angle of the animal to the speaker was systematically manipulated in 45° angles. With 0° being directly facing the speaker, we would reposition the animal at random 45° position offsets from forward in a random sequence for each recording session until all eight positions were completed. We broadcast 30 exemplars of noise stimuli with a 30 s inter-stimulus interval at each of the eight positions. We also measured subjects head position relative to the speaker in a subset of Freely-Moving and Communication contexts. For these test sessions, two cameras (GoPro Hero Session) recorded the animal’s position simultaneously from two locations. One was positioned to record the animal from the right side of the test box, while the other was positioned directly above the test box. During these sessions, an Arduino system flashed an LED visible to both cameras at 0.5 Hz. This signal was recorded along with the audio and neural streams as well to accurately align the video streams with the start of neural recording. Images at stimulus onset were captured for each session and the relative angle of the head to the speaker measured.

### Neurophysiological Recording Procedures

Subjects were surgically implanted with an acrylic head cap using previously described procedures (Courellis et al., 2019; Miller et al., 2015; Nummela et al., 2017). In a subsequent procedure, a 16ch Warp16 microelectrode array (Neuralynx) was implanted in prefrontal cortex. Each array had 16 independent guide tubes in a 4 × 4 mm grid with tungsten electrodes. Each array was implanted on the surface of the brain with each electrode in the guide tubes entering the laminar tissue perpendicularly when pushed by a Warp Drive pusher. The calibrated Warp Drive pusher would attach to the end of a guide tube to allow advancement of 10 to 20 μm per electrode twice a week.

Electrodes were recorded at 20,000 Hz with a prefilter at 1 Hz to 9000 Hz, and 20,000 gain. Subjects had a 1:1 gain headstage preamplifier connected to the Warp16 arrays that was attached to a tether to allow subjects to freely move around in the test box. A metal coil tightly wrapped around the tether prevented any interference by the subject during Freely and Communication contexts. Offline spike sorting was completed by combining across multiple sessions recorded in a single day, applying a 300 to 9000 Hz filter and thresholding subsamples across the entire recording session. After base thresholding, unit 1 ms waveforms were plotted in PCA space across the first three principal components and time. DBSCAN was used to automatically cluster the units followed by manual curations of all units to ensure proper clustering. Units with 13 dB SNR or greater and <1% violation of the 1 ms refractory period for inter-spike intervals were included in our analyses. Overall, 388 isolated single units were identified that met or exceeded these thresholds, with some channels collecting multiple well isolated single units. Of the 388 single units, 247 units met these thresholds for all three test contexts in a single session. Typically, neurons that failed to meet these criteria in all contexts did so because of noise introduced into the recording for one of the contexts.

### Perfusion, Tissue Processing and Reconstructions

At the conclusion of the study, animals were anesthetized with ketamine, euthanized with pentobarbital sodium, and perfused transcardially with phosphate-buffered heparin solution followed by 4% paraformaldehyde. The brain was impregnated with 30% phosphate-buffered sucrose and blocked. The frontal cortex was cut at 40 µm in the coronal plane. Alternating sections were processed for cytochrome oxidase (Wong-Riley, 1979), nissl substance with thionin, and vesicular glutamate transporter 2 (vGluT2). Areas were determined using previously identified criteria (Paxinos et al., 2012). Electrode penetrations, and tracer injection sites were used to reconstruct the location of the electrode array with respect to the anatomical borders and confirm the location of electrode penetrations. Images of the tissue were acquired using a Nikon eclipse 80i. These methods are similar to those employed in previous anatomical studies of marmoset neocortex (de la Mothe et al., 2006, 2012) and our earlier neurophysiology experiments (Miller et al., 2015; Nummela et al., 2017).

### Data Analysis

#### Stimulus Response Significance

Each single unit was classified as exhibiting a significant response to an acoustic stimulus if the firing rate during one of the three following stimulus periods exhibited a statistically significant change in activity at α=0.05 significance level with a Sign Rank test relative to the 500 ms prior to stimulus onset: [1] the entire duration of the stimulus, [2] the peak 500 ms firing rate of the unit within the duration of the stimulus, or [3] the first or second pulse of the phee call stimulus. These latter two criteria were adopted because multi-pulsed phee calls have a long duration (∼2500-3000 ms) and the observation that although many single neurons did not exhibit a sustained change in firing rate for the duration of these vocal signals, the firing rate increased substantially for some period of the stimulus.

#### Array Channel Responsiveness

To determine whether specific areas of marmoset PFC had a higher concentration of acoustic responsive neurons, we performed the following analysis: we divided the total number of well-isolated single units for each electrode channel that exhibited a significant response to at least one stimulus by the total single units recorded from that electrode. Each electrode location position in PFC is shown in Figure 1B along with a color representation of that ratio responsiveness.

#### PSTH Normalization

Unit trials were binned at 100 ms intervals starting 1000 ms prior to stimulus onset and ending 1500 ms after stimulus onset for Noise, Twitter, and Reverse Twitter. Phee and Reverse Phees included 4000 ms after stimulus offset. The firing rate for each bin within each trial for a given unit was calculated. Normalization occurred by calculating the mean firing rate and standard deviation of all the bins prior to stimulus onset to Z-score the firing rate for the bins following stimulus onset.

#### Firing Rate Normalization

To compare changes in firing rate between two contexts, we compared neural activity in the 1000 ms prior to stimulus onset with either the first pulse of the phee call or entire duration of the Noise stimulus. The first pulse and Noise duration were roughly the same duration on average: 1250 ms and 1000 ms, respectively. The firing rates prior to stimulus onset were used to Z-score the subsequent mean after onset Firing Rates. For analyses that only considered phee call comparisons, we used the entire duration of the phees and Z-scored in the same manner.

#### Spatial Selectivity Analysis

For the Orientation context outlined in the Test Procedures above, all 240 trials were analyzed using a 2-Way ANOVA. The firing rate in the 500 ms before and after onset were used for comparison across all trials, with trials organized into the 8 different 45° bins based on the relative angle of the subjects’ head to the speaker. If the unit was identified as stimulus responsive according to the metrics outlined in the Stimulus Response Significance section above, the interactive effect of spatial orientation was tested and Tukey-Kramer corrected comparisons were used to determine any orientation that had significance compared to the others. If all orientations exhibited similar responses with a main effect, we counted that unit as being generally responsive (all 8 orientations). In the Freely-Moving and Communication contexts, a similar analysis was used for Noise and Phee stimuli. As noted in Behavioral Recording Procedures, the gaze direction from the front of the head relative to the speaker on the transverse plane was marked. These orientations were binned into 8 different 45° groups similarly to the Orientation condition and the same analysis performed. The results from these analyses are shown in Figure S2A.

#### Inter-stimulus Interval Analysis

For each unit used in the analysis shown in Figure S2B, we calculated the firing rate of each trial in which a phee call was broadcast and the subsequent one with the same stimulus in each of the three behavioral contexts. The ratio of the firing rate for each of the two stimulus periods was calculated. Standard deviation was calculated by binning each 10 seconds and then finding the standard deviation 5 seconds before and after that bin time. 95% CI was calculated by the square root of the df of the calculated standard deviation divided by the χ^2^ inverse of that df. Only pairs of phee stimuli with an inter-stimulus interval less than 60 s were used in this analysis.

#### Linear Model

We conducted a linear mixed effect model (MATLAB function fitlme fit by maximum likelihood) to test for a linear effect of context (restrained < freely < communication) on normalized firing rate for all the maintained units. Context was coded as a numerical variable (restrained = 1, freely = 2, communication = 3) in order to test if it had a linear effect. The formula for the model was as follows using Wilkinson notation: FiringRate ∼ 1 + Context + (1 | Subject) + (1 + Context | Subject) + (1 | Channel:Session:Subject) + (1 | Session:Subject) + (1 | Type), where Type is the selectivity type of the neuron. The random effects in the model account for the nested design of the experiment where there were repeated observations for each channel within each session for each subject. The function anova using the Satterthwaite estimation of degrees of freedom was applied to the model to test for the significance of the effect of context.

#### Decoder Classification

Five hundred Monte-Carlo simulations were created for each model by subsampling each unit’s class of relevant trials with replacement. For single-unit decoders, 1000 samples were taken from each class as each class would have less than 100 samples per class. For multi-unit decoders that compared across multiple bins, we used 2161 samples calculated by multiplying and rounding the total number of possible units to include, the number of data points for each unit, and 1.25 to ensure enough representation. Equal numbers of samples were drawn for each class for each unit after 50% partition between training and test sets. Each simulation created new partitions for each unit. Each trial had 7 points of data (3 bins for each of the 2 pulses, and 1 bin for the inter-pulse-interval). The Samples × 7 matrix for each unit would then be combined with all other units with sufficient trials, for each of the classes within both training and test. For example, the 26 RFC units for Restrained, Freely, and Communication classes would produce a matrix of size 6483 × 182 for both training and test with no overlap of trials across those two. Each row is then assigned a value representing the class that all 182 values represent. PCA plots had difficulty representing the classes, so the MATLAB function tsne (t-distributed stochastic neighbor embedding) was used to represent the separability of classes. Only units with at least five events from each class of interest were included in these analyses. State-based classification used only four points of data as previously done (Nummela et al., 2017) rather than the seven points applied here. These points were the firing rate for the duration of the first pulse and second pulse, as well as 1.5 s before stimulus onset and 1.5 s after stimulus offset.

The training data was fit using the “fitcecoc” function in MATLAB using the default settings. This creates multiclass support vector machines for each simulation, and then predicts the class of the test data. Each confusion matrix that results from the predictions is then stored, and the mean value shown. We also quantified the performance of each classifier by the Matthews Correlation Coefficient (MCC). For a K × K confusion matrix C (i.e. K = 3 for Restrained, Freely, and Communication classes):

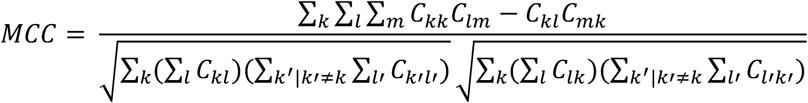

For K = 2, MCC ranges from -1 to +1. The +1 means perfect prediction, -1 means complete disagreement between predicted and actual, and 0 is no better than chance prediction. For K = 3, the lower limit is not at -1 and unique to any given classifier. Still, 0 means it is at chance prediction and +1 is perfect prediction. We conducted a null-hypothesis test for both classifiers running the same size data as the full data set (all 247 units). With a randomized assignment of class for each row in training and test data, the MCC was at 0 as expected.

## Figures

**Supplemental Figure 1.**
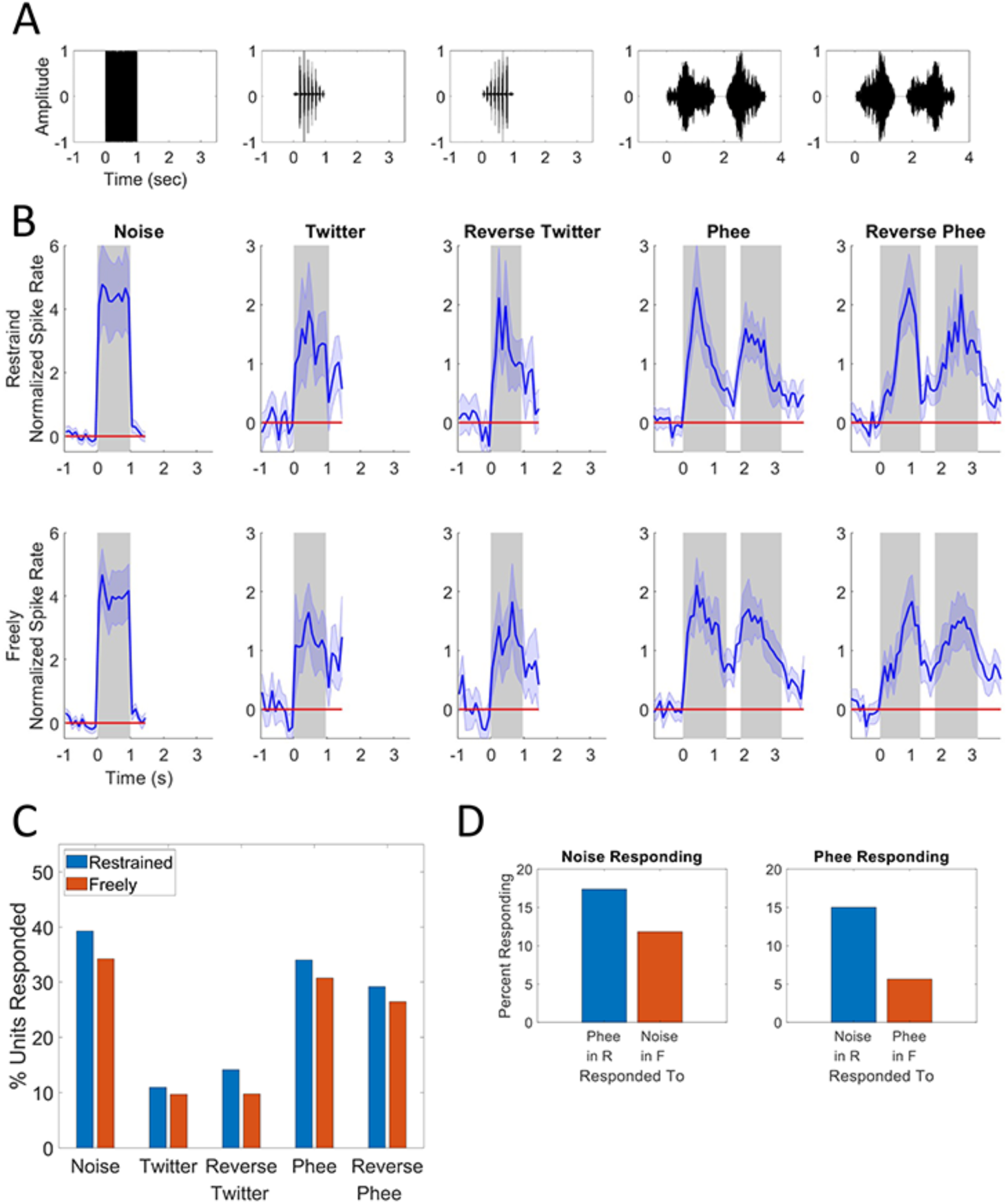
(A) Waveforms of the five stimulus sets presented. Left to right: noise, twitter, reverse twitter, phee, reverse phee. Same time scale and range is used to allow easier comparison. (B) Normalized PSTH plots for all neurons exhibiting a stimulus-driven response to each of the five stimulus types. Shaded areas represent 95% confidence interval while blue lines represent the mean firing rate. Gray boxes represent the full duration of a stimulus or the two pulses of the phee calls. Red line represents the baseline firing rate at standardized 0. Neural responses to the five stimuli in the Restrained (above) and Freely-Moving (below) are shown. Because noise, twitter, reversed twitter and reversed phee do not elicit responses from marmosets during natural communication, these stimuli were not presented in the Communication context. (C) Ratio of stimulus driven responses to a given stimulus category for all units found within each context. Restrained is in blue and Freely-moving in orange. (D) Probability of a single Noise responsive (left) and Phee responsive (right) neuron in the Restrained context exhibiting a significant stimulus-driven response to the opposite stimulus (Noise or Phee) in the Restrained context (blue) or the same stimulus in the opposite behavioral context (Restrained or Freely-Moving) is shown in orange. Similar to analyses of phee calls shown in Figure 2, neurons exhibited little cross-context reliability in their response, including for the same stimulus type.

**Supplemental Figure 2.**
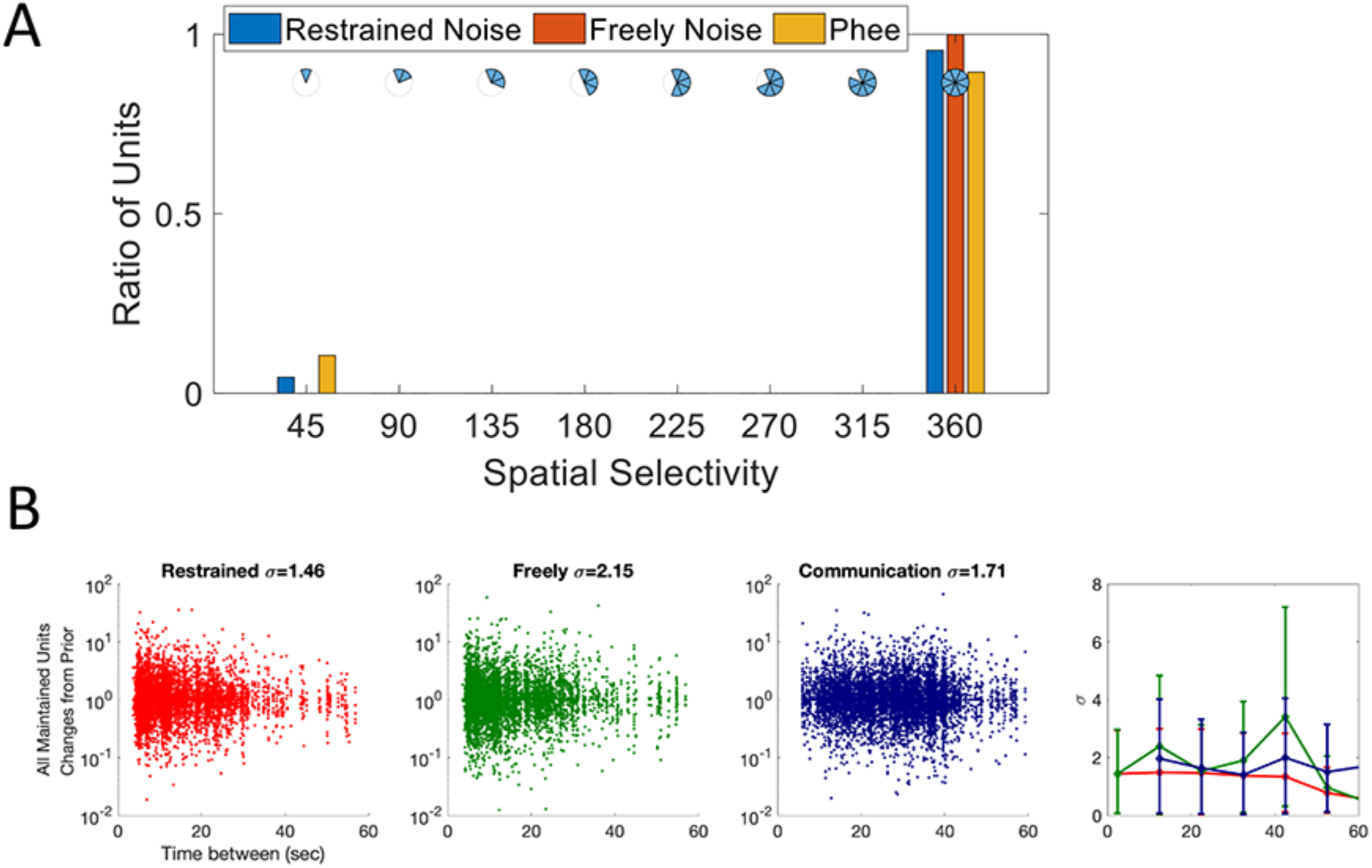
(A) Ratio of units exhibiting spatial selectivity of neural response. 45° indicates units that only exhibited a significant response when the relative head and speaker were at a single narrow direction, while 360° indicates neurons that exhibited the same response at all 8 directions tested. This includes tests for Noise in the Restrained (blue) and Freely-Moving (Orange) contexts, while Phee calls in the Freely-Moving and Communication contexts were combined. These analyses were performed on a subset of neurons: Restrained: Noise (n = 177 units); Freely-Moving: Noise (n = 103 units); Freely-Moving and Communication Phee calls (n = 94 units). (B) Trial-to-trial changes in firing rate for all 247 maintained units in the three contexts: Restrained (red), Freely-Moving (green) and Communication (blue). Log scale shows ratio of firing rate of second stimulus trial over first stimulus trial. 0/0 was set to 1. This was performed for all pairs of stimuli in which the inter-stimulus interval was <60 s. Each dot represents the firing rate ratio for each pair of consecutive stimuli. 95% confidence interval for each time bin’s standard deviation are shown on the right.

